# Interference of pH buffer with Pb^2+^-peripheral domain interactions: obstacle or opportunity?

**DOI:** 10.1101/2019.12.31.892232

**Authors:** Sachin Katti, Tatyana I. Igumenova

## Abstract

Pb^2+^ is a xenobiotic metal ion that competes for Ca^2+^-binding sites in proteins. Using the peripheral Ca^2+^-sensing domains of Syt1, we show that the chelating pH buffer Bis-Tris enables identification and functional characterization of high-affinity Pb^2+^ sites that are likely to be targeted by bioavailable Pb^2+^.

**Significance to Metallomics:** Syt1, a key regulator of Ca^2+^-evoked neurotransmitter release, is a putative molecular target of Pb^2+^. We demonstrate that the use of a chelating pH buffer Bis-Tris enables identification of Ca^2+^-binding sites that would be most susceptible to Pb^2+^ attack in the cellular environment. In addition, experiments conducted in Bis-Tris revealed the differences between the membrane-binding responses of two Ca^2+^-sensing domains of Syt1, C2A and C2B. This work advances the understanding of how Pb^2+^ interacts with multipartite Ca^2+^-binding sites, and illustrates that conducting the experiments under both chelating and non-chelating conditions could provide valuable insight into the mechanism of metallosensory proteins.

---

Lead (Pb^2+^) is a xenobiotic heavy metal ion that shows acute as well as chronic systemic toxicity in the human body.^1^ No extent of Pb^2+^ exposure is considered “safe”, and if undetected, can cause irreversible neurological damage in young children.^2, 3^ Among several proposed modes of Pb^2+^ toxicity, the ability to mimic Ca^2+^ is particularly alarming, as it highlights the vulnerability of ubiquitous, multi-site Ca^2+^-binding proteins towards Pb^2+^ attack.^4^ Identification of physiologically relevant Pb^2+^-binding sites on these proteins is challenging because Pb^2+^ interactions can be both specific as well as opportunistic, and often exhibit affinities higher than those of native metal ions.^5^ Synaptotagmin 1 (Syt1), a key regulator of Ca^2+^-evoked neurotransmitters release^6^ and putative molecular target of Pb^2+^,^7^ provides a remarkable example of this challenge. In the non-chelating environment such as MES buffer, four out of total five Ca^2+^-coordinating sites on the tandem C2 domains of Syt1 (C2A and C2B, **Fig. 1A**) bind Pb^2+^ with affinities higher than Ca^2+^.^8^ Site 1 on each domain exhibits sub-micromolar Pb^2+^ affinity, while Site 2 is substantially weaker in comparison (**Fig. 1A, table**).

**Figure 1.**
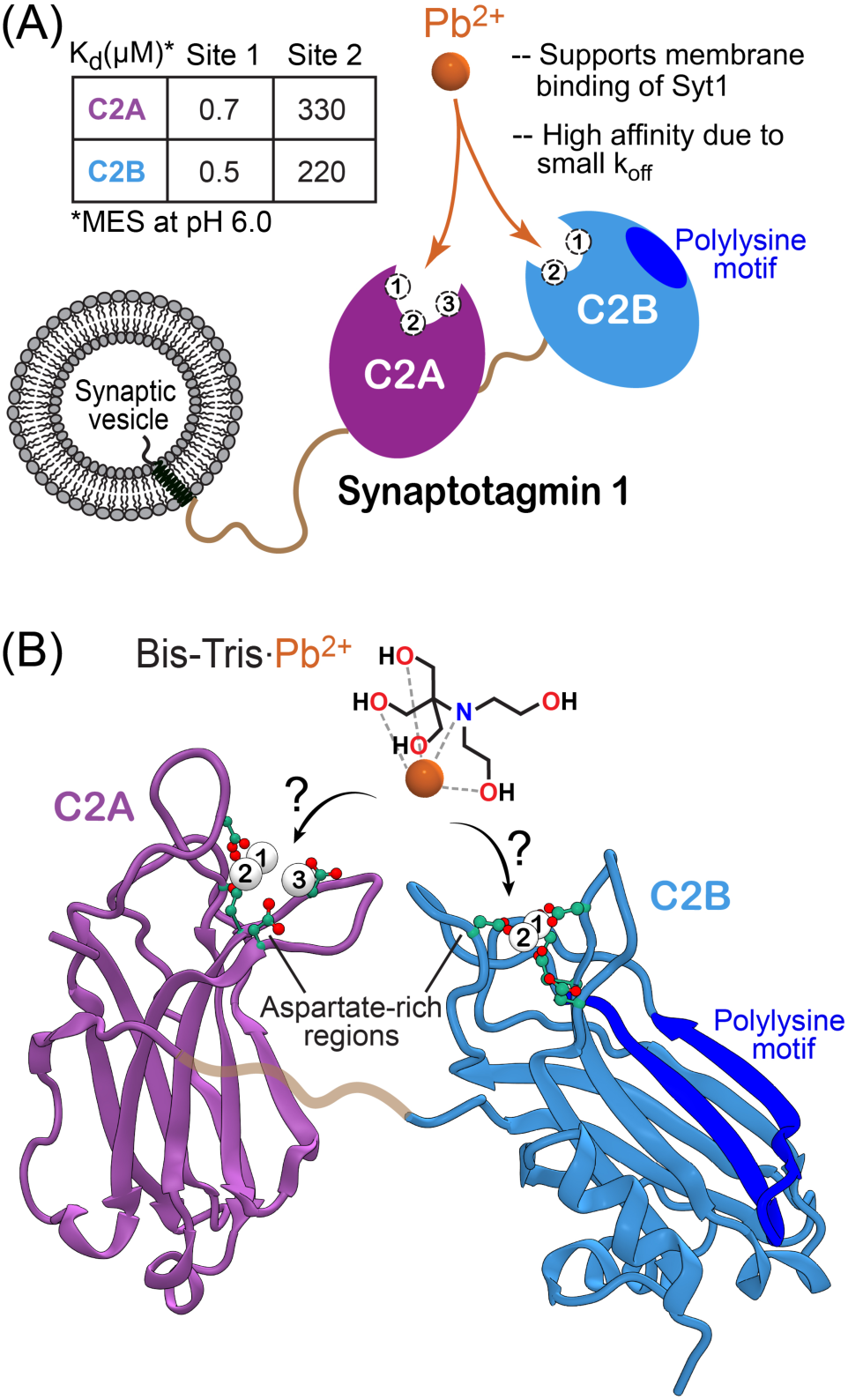
C2 domains of Syt1 interact with Pb^2+^ ions through the aspartate-rich loop regions. **(A)** Schematic representation of Syt1, a neuronal transmembrane protein and its cystosolic C2 domains, C2A and C2B. Ca^2+^-binding sites are labeled individually in each C2 domain. The affinities of Pb^2+^ to Sites 1 and 2 measured in MES buffer at pH 6.0 are given in the table.^8^ Pb^2+^ and Ca^2+^ do not appreciably bind to Site 3 under these conditions. **(B)** 3D structures of C2A (1BYN) and C2B (1UOW) domains showing the location of the Ca^2+^-binding aspartate-rich sites within the apical loop regions. Also shown is the schematic representation of the Pb^2+^ complex with Bis-Tris. The exact coordination geometry of Pb^2+^ in the Bis-Tris complex is unknown.

Here, we demonstrate, using the C2 domains of Syt1 as a paradigm, an approach that uses a pH buffering agent and Pb^2+^ chelator, Bis-Tris, to selectively probe high-affinity Pb^2+^ sites and determine their specific functional roles. Bis-Tris has five hydroxyl groups and one tertiary amine group (**Fig. 1B**), all of which could potentially serve as ligands for a wide array of divalent metal ions.^9^ While the challenges of using chelating pH buffers for metal ion-binding studies are known,^10^ Bis-Tris was our chelator of choice because, compared to other strong chelating agents such as EDTA, it forms weaker yet stable complexes with Pb^2+^.^9^ In addition, Bis-Tris is a highly adaptable chelator in that the number of its functional groups engaged in metal-ion coordination can vary depending on the presence of competing ligands donated, e.g., by proteins, lipids, or other small molecules. We reasoned that due to these properties, Bis-Tris would be a suitable mimic of a chelating physiological environment. In this work, we show that not only the use of Bis-Tris enables discrimination between high- and low-affinity Pb^2+^ sites, it also provides mechanistic information on the metal-ion dependent association of C2 domains with anionic membranes. We therefore argue that a judicious use of chelating pH buffers in metal-binding studies could provide a valuable insight into the mechanisms of metallosensory proteins.

To determine which of the five metal-binding sites of Syt1 are populated by Pb^2+^ in Bis-Tris, we conducted NMR-detected Pb^2+^-binding experiments on the individual C2 domains, C2A and C2B. Solution NMR spectroscopy is best suited for this purpose because it enables clear identification of metal-ion binding sites with a wide range of affinities, as demonstrated previously for the C2 domains of Syt1 and protein kinase C (PKC).^8, 11^ The information about protein-metal ion interactions is obtained by collecting 2D ^15^N-^1^H hetero-nuclear single-quantum coherence (HSQC) spectra of the uniformly ^15^N-enriched ([U-^15^N]) C2 domains, in the presence of different concentrations of metal ions. Binding of a metal ion alters the electronic environment of the backbone amide N-H_N_ groups that are proximal to the binding site, which results in chemical shift changes of the ^1^H_N_ and ^15^N nuclei. Given the tendency of certain buffering agents to interact with proteins,^12^ prior to Pb^2+^ addition, we compared the metal-free NMR spectra of the C2 domains in Bis-Tris and MES buffers. For C2A, the spectra in MES and Bis-Tris were identical (**Fig. S1A**). For C2B, a subset of N-H_N_ resonances showed small variations in their chemical shifts as well as intensities (**Fig. S1B**), indicating possible weak interactions with one of these buffering agents. Because of the highly basic nature of C2B and its propensity to interact with polyanionic molecules,^13^ the weakly interacting buffer is likely to be MES.

Upon addition of Pb^2+^ to the C2A domain at a molar ratio of 2:1 (C2A:Pb^2+^), we observed an appearance of a subset of N-H_N_ cross-peaks that belong to the Pb^2+^-complexed protein species (top inset of **Fig. 2A**, cyan). Adding more Pb^2+^ to achieve a 1:1 molar ratio results in the formation of a single protein species, the C2A·Pb1 complex (**Fig. 2A** and top inset, blue), where Pb^2+^ is bound to Site 1. This behavior is identical to that in the non-chelating MES buffer,^8^ suggesting that Bis-Tris does not interfere with Pb^2+^ binding to Site 1 of the C2A domain and full saturation of this site can be achieved at a stoichiometric ratio of protein to Pb^2+^.

**Figure 2.**
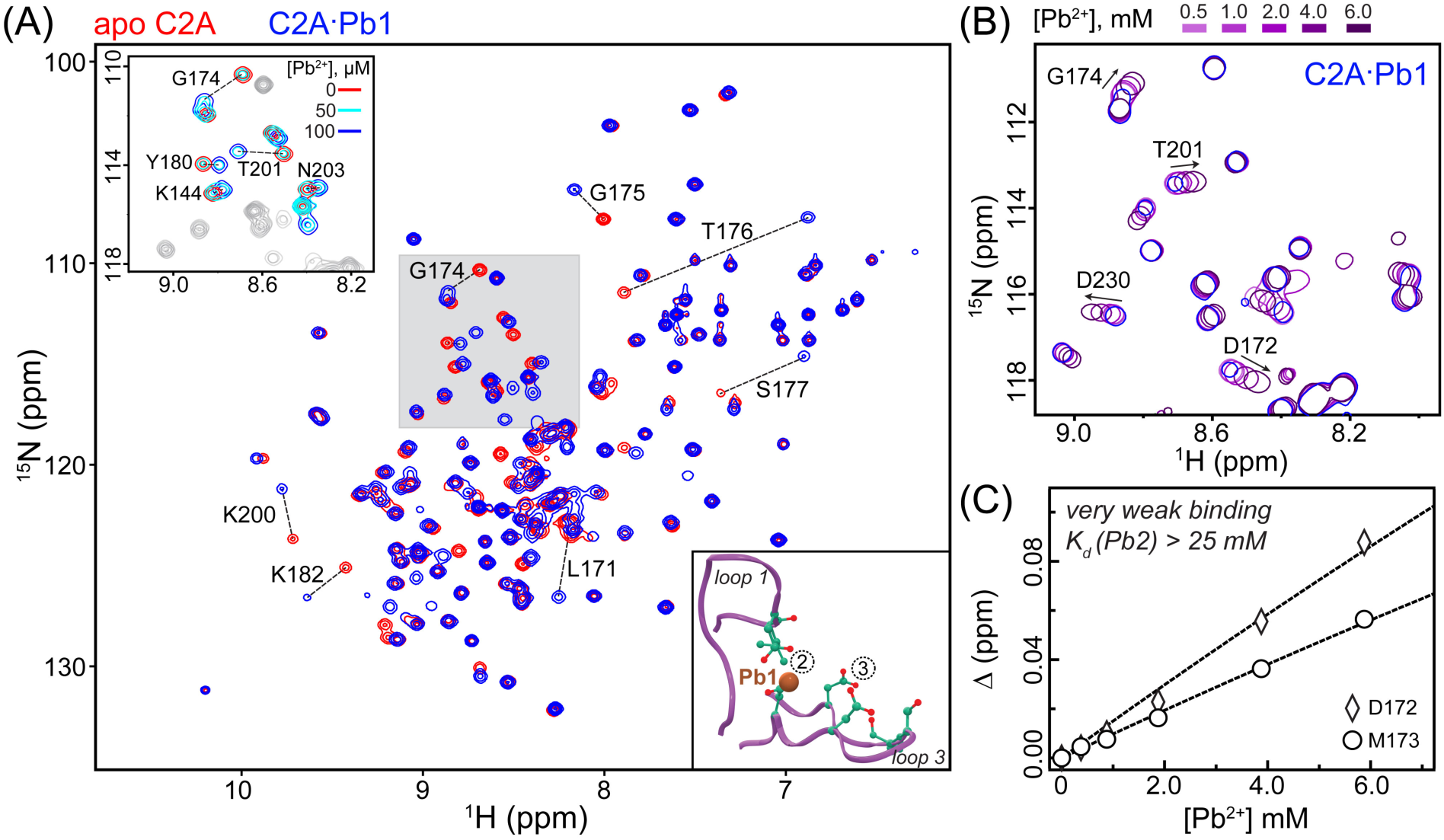
Bis-Tris inhibits Pb^2+^ binding to Site 2 but not Site 1 of the C2A domain. **(A)** Overlay of the [^15^N-^1^H] HSQC spectra in the absence (apo) and presence of stoichiometric Pb^2+^ (C2A:Pb^2+^ 1:1, 100 μM each) in 20 mM Bis-Tris buffer at pH 6.0. Residues showing response to population of Site 1 by Pb^2+^ are labeled. The spectra were collected at 25 °C on the Avance III NMR instrument (Bruker Biospin) operating at the magnetic field of 14.1 T and equipped with a cryogenically cooled probe. Top inset: expansion of the shaded spectral region with an additional Pb^2+^ concentration point (50 μM, C2A:Pb^2+^ 2:1, cyan) to illustrate the distinct chemical shifts of apo C2A and the C2A·Pb1 complex. Bottom inset: loop regions of the C2A domain showing the positions of Sites 1-3. **(B)** Overlay of the [^15^N-^1^H] HSQC expansions showing the chemical shift perturbations of several residues in the C2A·Pb1 complex due to Pb^2+^ binding to Site 2. **(C)** Chemical shift perturbations Δ (calculated as described in ref. 8) of two representative residues plotted as a function of Pb^2+^ concentration. The linear, non-saturatable profiles indicate that binding is extremely weak with a K_d_ of >25 mM. The lines are to guide the eye.

We were surprised to find however that addition of more Pb^2+^, with the objective to saturate Site 2 of the C2A domain, did not produce a drastic change in the N-H_N_ cross-peak chemical shifts (**Fig. 2B**) that we previously observed in MES.^8^ The Pb2 binding event is in the “fast” exchange regime on the NMR chemical shift timescale, where a single population-weighted cross-peak for a given residue shows a smooth trajectory in response to increasing Pb^2+^ concentration. This behavior enabled us to construct a binding curve by plotting the chemical shift perturbation Δ versus Pb^2+^ concentration. The curve is linear, with no indication of reaching the cusp region even at 60-fold excess of Pb^2+^ to protein (**Fig 2C)**. For comparison, in MES, Pb^2+^ was able to saturate Site 2 of the C2A domain at ∼25-fold excess to produce a K_d_ of 330 μM (see **Fig. 1A**, table).^8^ We observed the exact same pattern of Pb^2+^ interactions with C2B in Bis-Tris: complete population of Site 1 at the stoichiometric protein-to-Pb^2+^ ratio, and extremely weak binding to Site 2 (**Fig. S2**). We conclude that Bis-Tris does not affect Pb^2+^ binding to the high-affinity Site 1 but inhibits the interactions of Pb^2+^ with Site 2 of both C2 domains of Syt1. The affinity of Pb^2+^ to Site 3 of the C2A domain is too weak to be appreciably populated in either buffer.

We then asked if the Pb^2+^ dependent membrane-binding function of the C2 domains is influenced by Bis-Tris. When Pb^2+^ populates high- as well as low-affinity sites, as is the case in the MES buffer, both C2 domains interact readily with anionic membranes.^8^ However, when we conducted the C2A-vesicle co-sedimentation experiments in Bis-Tris, we found that Pb^2+^ was unable to support the membrane-binding function of the C2A domain. The fraction of the Pb^2+^-complexed protein bound to PtdSer-containing large unilamellar vesicles (LUVs) does not exceed 5%, even under conditions of the 40-fold molar excess of Pb^2+^ over C2A (**Fig. 3A**). A control experiment with the native metal ion, Ca^2+^, produced a fractional population of the membrane-bound species comparable to that observed in MES.^8^ Clearly, the effect of Bis-Tris is specific to the Pb^2+^-C2A-anionic membrane system. To ascertain that this finding is not technique-dependent, we conducted FRET-detected protein-to-membrane binding experiments shown schematically in **Fig. 3B**. In the presence of Ca^2+^, the FRET curve of the C2A domain represents a typical membrane binding response reported for the C2 domains (**Fig. 3C**).^11, 14^ In contrast, we observed no Pb^2+^-driven interactions of C2A with membranes in Bis-Tris, which is fully consistent with the co-sedimentation data of **Fig. 3A**. We therefore conclude that Bis-Tris interferes with the membrane-binding function of the Pb^2+^-complexed C2A domain.

**Figure 3.**
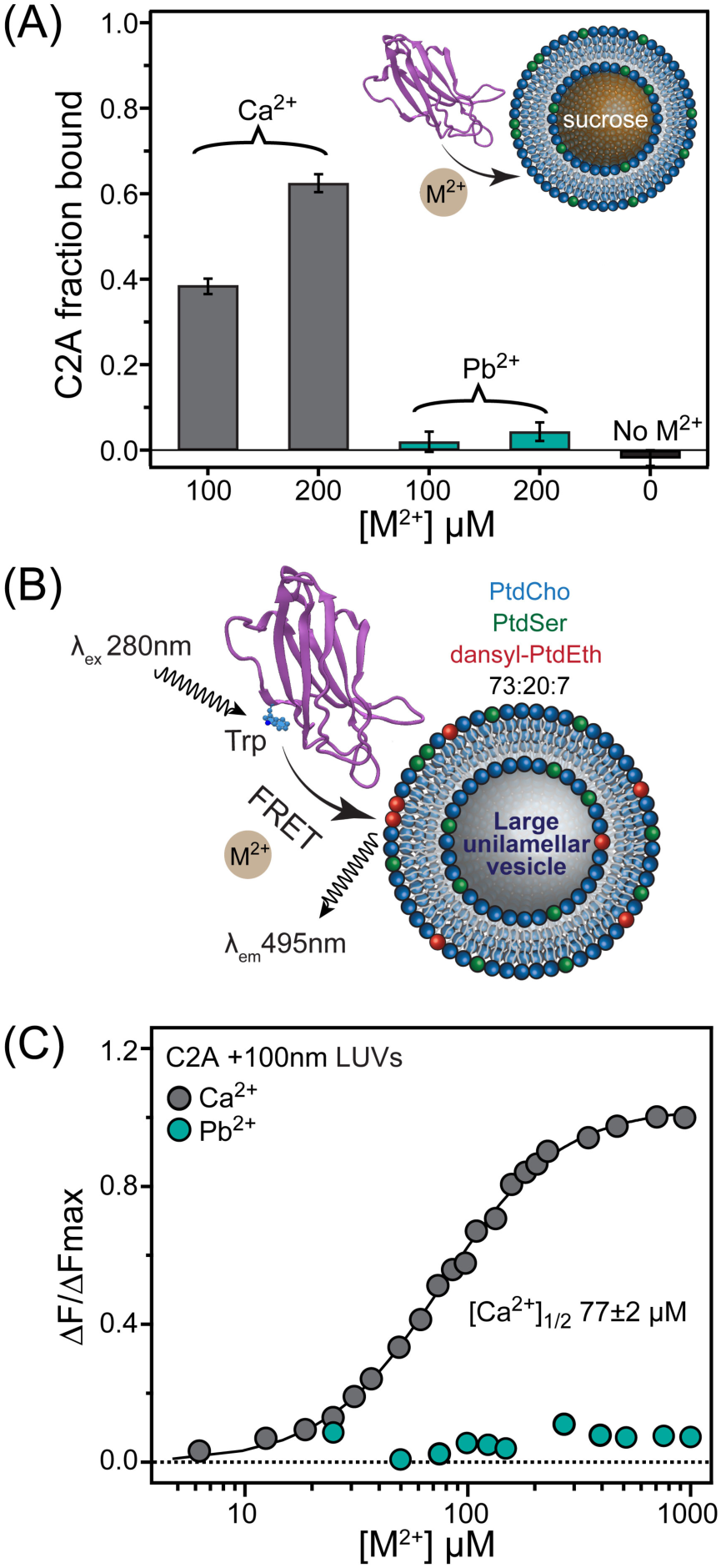
Pb^2+^-dependent membrane binding of C2A is abolished in the presence of Bis-Tris. **(A)** Fraction of C2A bound to sucrose-loaded LUVs in the presence of Ca^2+^ and Pb^2+^. The co-sedimentation experiments were conducted in 20 mM Bis-Tris at pH 7.0 and 150 mM KCl. The LUVs were 100 nm in diameter and consisted of PtdCho:PtdSer=80:20 (%, molar). The C2A concentration was 5 μM. **(B)** Schematic representation of FRET-detected C2-membrane binding experiments. The resonance energy transfer takes place between native Trp residues and dansyl-PE fluorophore embedded into LUVs. **(C)** FRET-detected C2A-membrane binding curves plotted as a function of increasing M^2+^ (M=Ca, Pb) concentrations. The change in fluorescence intensity at 495 nm, ΔF, was normalized to the maximum change observed in Ca^2+^-dependent experiments, ΔF_max_. Same buffer as for co-sedimentation experiments was used. The C2A concentration was 0.5 μM.

To understand the underlying cause of this behaviour, we considered two factors that are essential for driving the C2A-membrane interactions:^15^ (1) the general “electrostatic shift” due to metal-ion binding, which alters the charge of the intra-loop region from negative to positive; and the specific interactions of anionic phospholipid headgroups with the C2-complexed metal ions (and potentially the basic residues of C2A surrounding the loop region). Our data indicate that inhibition of Pb^2+^ interactions with Site 2 by Bis-Tris locks the C2A domain in the single Pb^2+^-bound state, and that the resulting electrostatic shift is insufficient for the C2A to engage in high-affinity interactions with anionic membranes.

We next sought to investigate if specific interactions of the Pb^2+^-complexed C2A domain with PtdSer are influenced by Bis-Tris. In MES, the C2A·Pb1 complex prepared by mixing stoichiometric amounts of C2A and Pb^2+^ associates weakly with PtdSer-containing bicelles.^16^ The chemical exchange process between the C2A·Pb1-PtdSer ternary complex and the C2A·Pb1 binary complexes occurs on the “fast-to-intermediate” NMR chemical shift timescale. This produces a specific pattern of resonance intensity decrease in the NMR spectra that predominantly affects the loop regions of the C2A domain. Here, we used a short-chain PtdSer analogue, 1,2-dihexanoyl-sn-glycero-3-[phospho-L-serine] (DPS) to determine how Bis-Tris influences the interactions of the C2A·Pb1 complex with PtdSer. In MES, addition of 5 mM DPS to 100 μM [U-^15^N enriched] C2A·Pb1 complex resulted in attenuation of the cross-peak intensities. The attenuation was quantified as the ratio of the N-H_N_ cross-peak intensities in the absence (I_0_) and presence (I) of DPS. Consistent with previous data obtained using PtdSer containing bicelles,^16^ residues that belong to the loop regions showed significant decrease in I/I_0_ values, along with several residues in the polylysine region between loops 1 and 2 (**Fig. 4A**). In contrast, the C2A·Pb1 complex in Bis-Tris showed little attenuation of signals that belong to the loop regions (**Fig. 4B**). This is especially evident in the difference I/I_0_ plot (**Fig. 4C**) that was constructed by subtracting the data in (**B**) from the data in (**A**). We conclude that specific interactions of the C2A·Pb1 complex with PtdSer, while clearly present in the non-chelating MES buffer, are absent in the chelating Bis-Tris buffer. How does Bis-Tris inhibit the interactions of the C2A·Pb1 complex with PtdSer? There are examples of crystal structures of protein-metal ion complexes where Bis-Tris provides additional ligands to the protein-bound metal ions.^17-19^ It is therefore plausible that Bis-Tris directly coordinates Pb1 and thereby prevents Pb^2+^ to form coordination bonds with the oxygens of PtdSer. This scenario is schematically shown in **Fig. 4D**.

**Figure 4.**
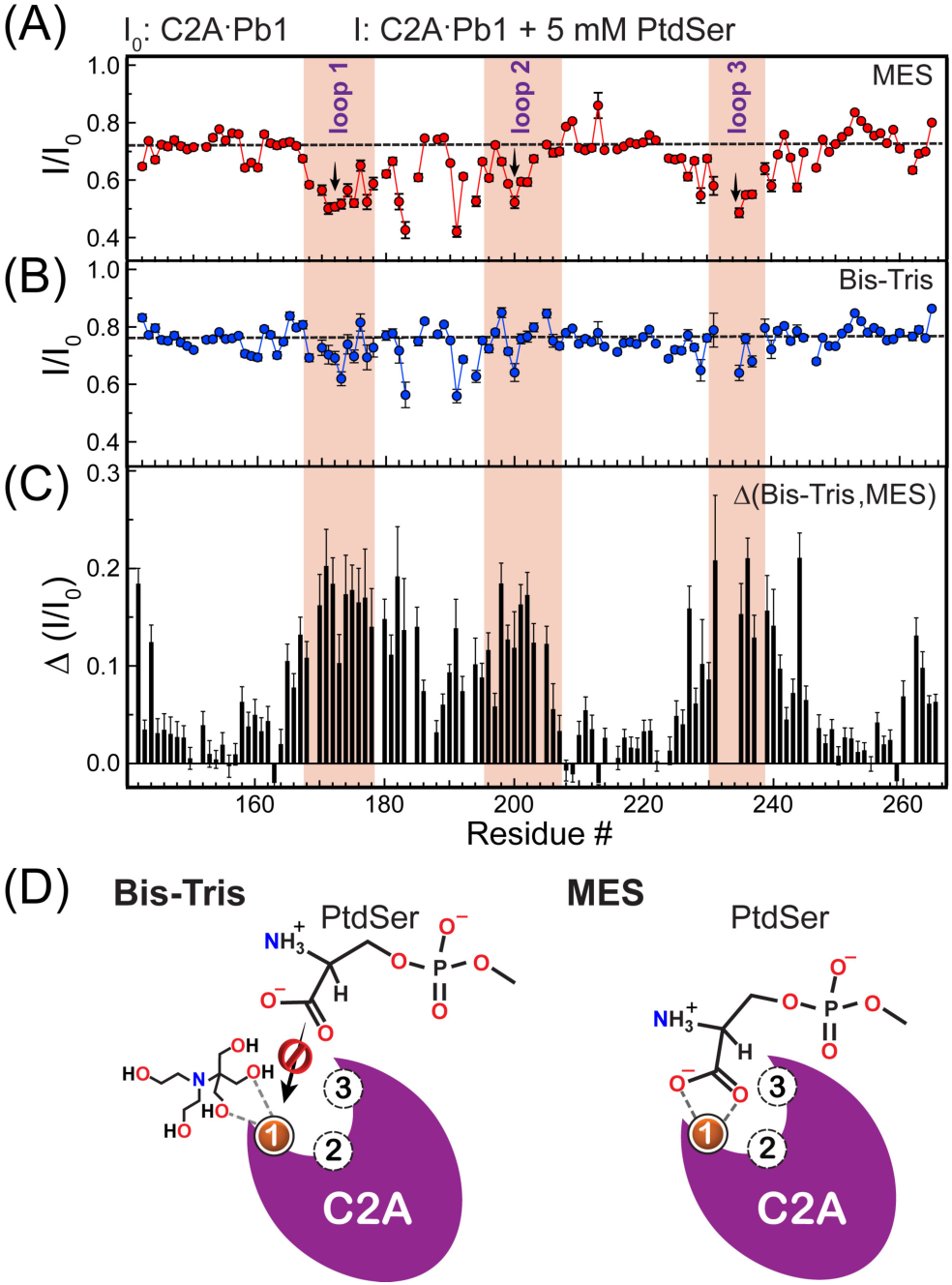
Pb^2+^-complexed C2A domain does not appreciably interact with PtdSer in the presence of Bis-Tris. Attenuation of peak intensities in the C2A·Pb1 complex upon addition of 5 mM DPS, expressed as residue-specific I/I_0_ ratios, where I and I_0_ are the peak intensities in the presence and absence of PtdSer, respectively. The data were collected at pH 6.0 in 20 mM **(A)** MES and **(B)** Bis-Tris buffers. The errors in the intensity ratios were estimated using the r.m.s.d. of the base-plane noise in each of the collected spectrum. **(C)** The difference between I/I_0_ ratios, demonstrating that the specific interaction of the C2A·Pb1 loops with PtdSer is present in the MES but not in the Bis-Tris buffers. **(D)** Schematic model showing how Bis-Tris can potentially interfere with C2A·Pb1-PtdSer interactions.

In conclusion, Bis-Tris eliminates two factors that contribute to metal ion-driven C2A-membrane interactions: the effect of general electrostatics, by inhibiting Pb^2+^ binding to Site 2; and the effect of C2A·Pb1-PtdSer interactions, likely by completing the coordination sphere of C2A-bound Pb1. These findings provide the following insight into the metal-ion dependent membrane binding function of the C2 domains. C2 domains with partially occupied metal-ion binding sites, through weak association with anionic membranes, are brought close to the polar membrane region.^16, 20^ It is believed that the resulting proximity to anionic lipid moieties elicits cooperative metal-ion binding to the remaining weaker sites by enhancing their affinities. This mutual cooperativity enables the domain to perform its function at physiological concentrations of Ca^2+^.^14^ Our results support this mechanistic model: by interfering with the PtdSer interactions of the C2A·Pb1 complex, Bis-Tris also precludes the cooperative enhancement of Site 2 affinity towards Pb^2+^. As a result, the domain stays locked in Pb1-bound state, unable to achieve complete electrostatic shift required for membrane association.

When viewed from the perspective of Pb^2+^ toxicity, our data indicate that in the cellular environment where Bis-Tris-like chelators and metabolites are abundant,^21^ the membrane interactions of the C2A·Pb1 complex are unlikely to occur. Combined with our previous finding that the presence of Pb^2+^ in Site 1 of C2 domains desensitizes them to further Ca^2+^ binding,^8, 11^ the ability of Pb^2+^-complexed C2A domain to interact with membranes is unlikely to be restored or rescued at physiological Ca^2+^ concentrations.

The next step was to determine how Pb^2+^-complexed C2B interacts with anionic membranes in Bis-Tris. The C2B domain is significantly more basic than C2A due to the presence of a distinct polylysine motif (see **Fig. 1B**), and additional positively charged residues, Arg 398 and Arg 399, at the end opposite to the loop regions. Although the polylysine region is located several Ångstroms away from the metal-ion binding loops, it plays a critical role in the membrane association process.^22-24^ In particular, its interaction with the second messenger PtdIns(4,5)P_2_ dramatically increases the Ca^2+^ affinity to C2B, lowering the Ca^2+^ concentration threshold required for the membrane association of this domain.^25^ The underlying basis of this cooperative effect is the change of the electrostatic potential of C2B, caused by the interactions of its basic regions with anionic phospholipids. The cooperativity between anionic phospholipids and divalent metal ions is mutual: binding of metal ions to the loop regions enhances the interactions of C2B with phospholipids.

In stark contrast to C2A, Pb^2+^-complexed C2B associates with LUVs in Bis-Tris (**Fig. 5A**, compare with **Fig. 4A**), with fractional population of the membrane-bound protein species comparable to that observed in presence of Ca^2+^. The FRET experiments were carried out by titrating the PtdSer-containing LUVs into a solution containing C2B and excess Pb^2+^, to minimize C2B-induced LUV clustering and associated increase in scattering.^26^ The FRET results mirrored those of the co-sedimentation experiments. Pb^2+^ was almost as effective as Ca^2+^ in driving protein-membrane association, producing [PtdSer]_1/2_ values of 3.4 μM compared to 2.8 μM, respectively (**Fig. 5B**). The two plausible explanations for this behaviour are: (1) the inherently basic nature of the C2B surface that enables membrane interactions in single Pb^2+^-bound state; and (2) the allosteric enhancement of C2B affinity to Pb^2+^ upon interactions of PtdSer with basic regions of the protein.

**Figure 5.**
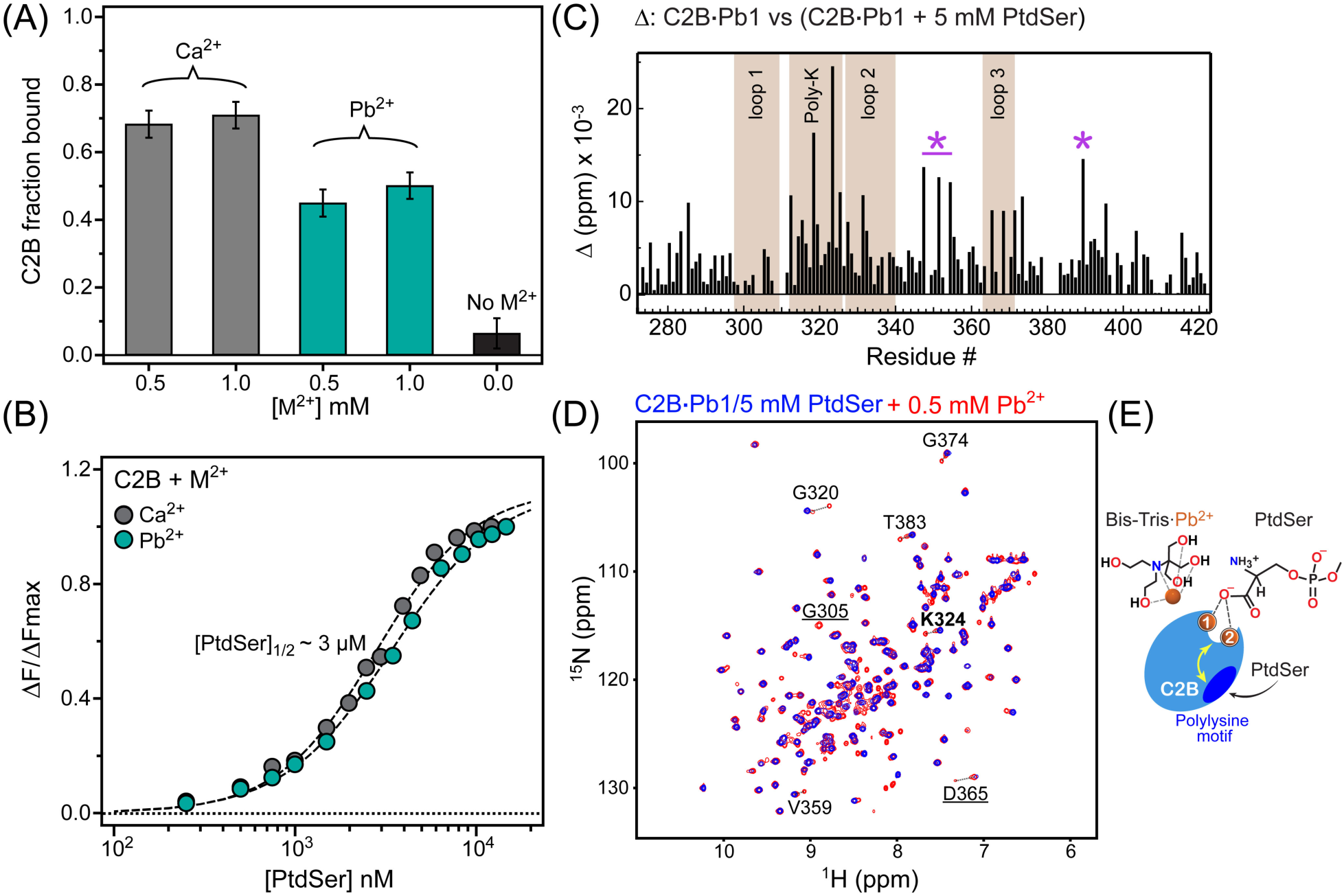
Pb^2+^-dependent membrane binding of C2B persists in the presence of Bis-Tris. **(A)** Vesicle co-sedimentation experiments conducted in 20 mM Bis-Tris and 150 mM KCl at pH 7.0 show that in contrast to C2A, Pb^2+^-complexed C2B associates with anionic membranes. The C2B concentration is 5 μM. **(B)** C2B-to-membrane binding curves plotted as normalized FRET efficiency versus accessible PtdSer concentration in LUVs. The concentrations of C2B and M^2+^ are 0.5 μM and 500 μM, respectively. Pb^2+^ is almost as effective as Ca^2+^ in driving membrane interactions. **(C)** Chemical shift perturbations Δ caused by addition of DPS (cmc ∼ 12 mM) to the C2B·Pb1 complex are plotted as a function of primary structure. The polylysine motif and residues at the “bottom” of C2B, marked by asterisks, are the likely interaction sites based on the chemical shift data. **(D)** Overlay of the [^15^N-^1^H] HSQC spectra showing an additional set of cross-peaks that appears upon addition of 0.5 mM Pb^2+^ to the C2B·Pb1 complex in the presence of 5 mM DPS. The newly formed Pb^2+^-bound C2B species are in slow exchange with the C2B·Pb1 species, as illustrated for residues G320, G374, T383, D365, and V359. The underlined residues G305 (loop 1) and D365 (loop 3) are broadened beyond detection in C2B·Pb1 but re-appear in the spectra upon Site 2 population by Pb^2+^. K324, shown in boldface, is a residue that belongs to the polylysine motif. **(E)** A model that schematically illustrates how interactions of anionic phospholipids with the polylysine motif of C2B could cause cooperative enhancement in Pb^2+^ affinities for the metal-ion binding sites of the C2B domain. This effect would enable C2B to effectively compete with Bis-Tris for Pb^2+^ and acquire a full complement of Pb^2+^ ions needed for membrane interactions.

To assess the relative contribution of these factors, we added DPS to the [U-^15^N] C2B·Pb1 complex (prepared by mixing stoichiometric amount of C2B and Pb^2+^) and compared the chemical shifts of the backbone amide groups between the NMR spectra of DPS-free and DPS-containing samples of C2B·Pb1. We observed modest changes in the chemical shifts of the C2B·Pb1 resonances that are located predominantly near the polylysine motif and the adjacent “bottom” part of C2B (**Fig. 5C**), suggesting that these regions are the primary PtdSer interaction sites. The nature of the C2B·Pb1-DPS interactions is electrostatic and is therefore expected to produce large chemical shift changes if the protein is fully DPS-bound. The small values of chemical shift perturbations suggest that the fractional population of the DPS-bound C2B·Pb1 species is small. Therefore, the inherently basic nature of C2B alone cannot explain the affinity of Pb^2+^-complexed C2B species to anionic membranes in Bis-Tris.

Next, we tested if the presence of PtdSer in solution enhances the affinity of C2B towards Pb^2+^ to the extent that the domain acquires an ability to compete with Bis-Tris and bind the second Pb^2+^ ion. Upon addition of 5-fold molar excess of Pb^2+^ to the C2B·Pb1 in the presence of 5 mM DPS, a new subset of cross-peaks appeared in the NMR spectrum (**Fig. 5D**, red spectrum). This indicates the formation of new Pb^2+^-bound species of the C2B domain that are in slow exchange on the NMR chemical shift timescale with the C2B·Pb1 complex. The only possible Pb^2+^ interaction site with the C2B domain is Site 2. Indeed, residues whose cross-peaks are typically exchange-broadened in the C2B·Pb1 complex but only appear upon population of Site 2, G305 and D365, are detectable in the sample containing Pb^2+^ excess. For several residues, more than two sets of resonances could be seen, reflecting the complex speciation of Pb^2+^-bound C2B in the presence of DPS. In the absence of PtdSer, the population of Site 2 in C2B by Pb^2+^ is negligible even at 60-fold molar Pb^2+^ excess (**Fig. S2**). Therefore, we conclude that neutralization of the polylysine region via interactions with PtdSer and, to generalize, anionic phospholipids, is the dominant factor that enables the weaker Site 2 of C2B to successfully compete with Bis-Tris for Pb^2+^. The resulting acquisition of a full complement of Pb^2+^ ions by the C2B domain, and the mutual cooperativity between the polylysine motif and metal-ion binding loop regions drives the membrane association of C2B. This scenario is schematically illustrated in **Fig. 5E**. In contrast to C2B, C2A showed barely any response to the addition of excess Pb^2+^ in the presence of 5 mM DPS (**Fig S3**).

In full-length Syt1, the C2A and C2B domains are connected by a flexible 9-residue linker and together form the full Ca^2+^-sensing unit (**Fig. 1B**).^27^ To determine how Pb^2+^-driven interactions of the C2AB fragment depend on the chelating properties of the environment, we conducted FRET-detected protein-to-membrane binding experiments in MES and Bis-Tris (**Fig. 6A**). Due to the multivalent membrane binding mode of C2B^28^ and the ability of the tandem C2 domains to interact with LUVs in “trans” mode with respect to each other,^29^ the C2AB fragment shows significant clustering of LUVs. The clustering thresholds are marked in red (**Fig. 6A**), and correspond to the accessible PtdSer concentration upon exceeding which visible precipitation of LUVs and concomitant decrease of FRET signal occur.

**Figure 6.**
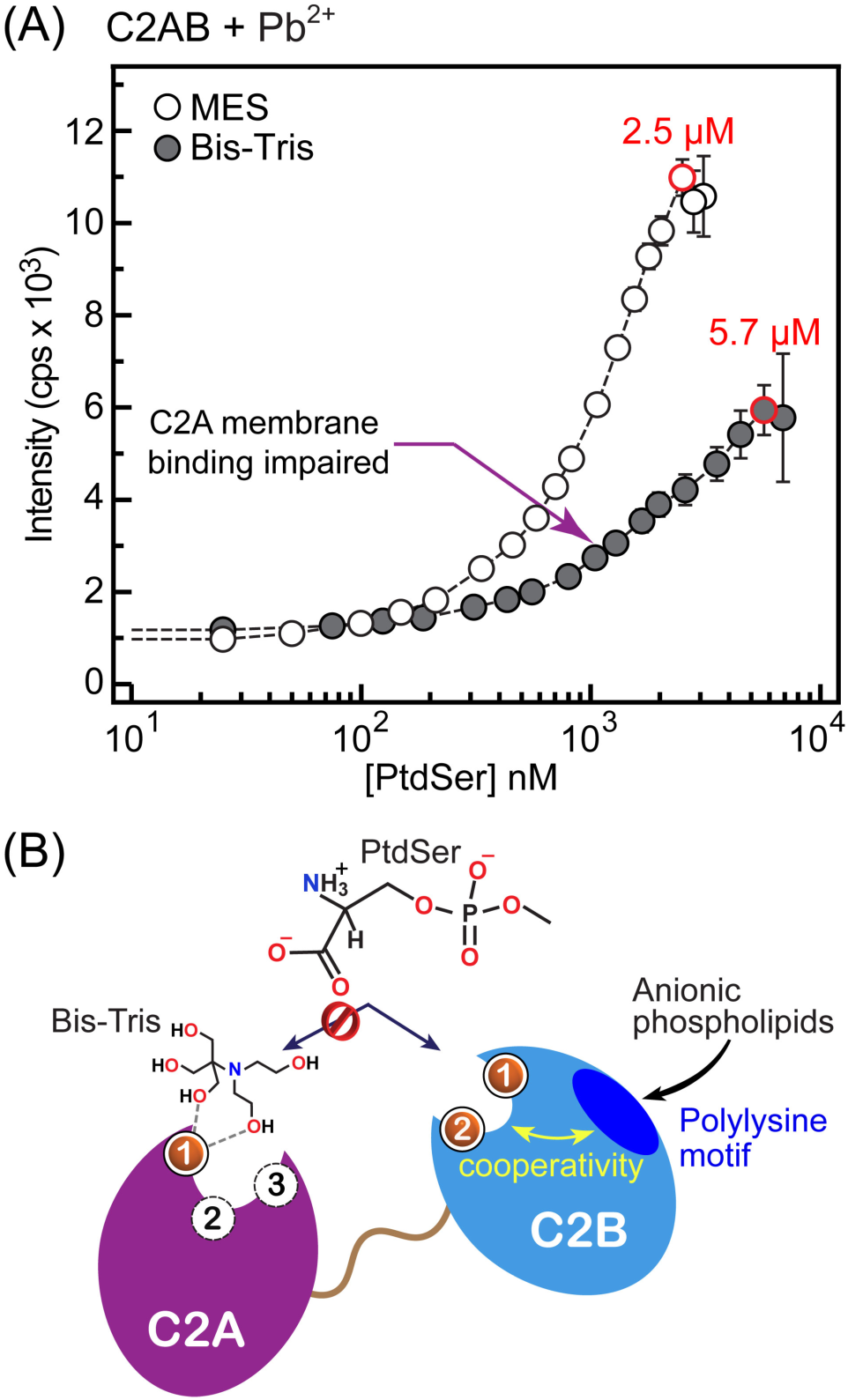
Membrane association of C2AB reflects the differential effect of Bis-Tris on the individual C2 domains. **(A)** C2AB-to-membrane binding curves plotted as a function of FRET efficiency versus concentration of accessible PtdSer in 100 nm LUVs. The concentration of C2AB and Pb^2+^ are 0.5 μM and 500 μM, respectively. The composition of LUVs is PtdSer:PtdCho:dansyl-PE=73:20:7 (molar). The buffer conditions are 20 mM Bis-Tris (MES) at pH 7.0 (6.0) with 150 mM KCl. The clustering point, at which light scattering by LUVs precludes further FRET measurements, is marked in red. More than 50% intensity difference in the dansyl-PE fluorescence intensity at the clustering point is observed in Bis-Tris compared to MES. **(B)** Schematic representation of the C2AB-Pb^2+^-anionic phospholipid interactions in the Bis-Tris environment. The association of anionic phospholipids with the polylysine cluster enables the population of Site 2 by Pb^2+^ and the coordination of protein-bound Pb^2+^ ions by PtdSer. Association of C2A with anionic membranes is extremely weak, because Bis-Tris interferes with Pb^2+^ binding to Sites 2 and 3, and PtdSer coordination by Pb1.

The most drastic difference between the binding curves of **Fig. 6A** is the overall decrease of FRET efficiency in the Bis-Tris buffer: at 2.5 μM accessible PtdSer, the FRET efficiency is ∼3-fold lower than that in MES. This suggests that C2AB-membrane interactions are significantly weakened in the presence of Bis-Tris. Based on the properties of individual domains in Bis-Tris (**Figs. 3** and **5**), this behaviour is likely caused by the impairment of the C2A membrane-binding function. This conclusion is further supported by the observation that the clustering threshold in Bis-Tris requires more than twice the accessible PtdSer, 5.7 μM, than that in MES.

In addition to providing mechanistic information about the determinants of C2-membrane association, our data on the isolated domains and the C2AB fragment in Bis-Tris led us to propose a model of how Pb^2+^ would interact with Syt1 under the chelating conditions of the cellular milieu (**Fig. 6B**). Due to the competition with the physiological chelators such as glutathione and metallothionein,^21, 30^ only high-affinity Pb^2+^ binding sites of Syt1 will be populated, i.e. Site 1 of each C2 domain. Upon population of Site 1 by Pb^2+^, C2B domain will interact with anionic lipids of the presynaptic membranes, PtdIns(4,5)P_2_ and PtdSer,^16^ via the polylysine motif; due to the positive cooperativity, the affinity of C2B to divalent metal ions will be enhanced. As we demonstrated previously,^8, 11^ Ca^2+^ is unable to share ligands with Pb1 and occupy the remaining vacant metal ion sites. The reason is high electronegativity of Pb^2+31^ that results in protein-bound Pb1 depleting the electron density of the oxygen ligands^32^ shared by Sites 1 and 2. It is therefore feasible that instead of Ca^2+^, C2B·Pb1 will preferentially acquire Pb^2+^ at Site 2, and that will drive the membrane association of Syt1 via the C2B domain. Our results suggest that contribution of C2A to membrane binding in the chelating environment will be significantly attenuated, because the C2A·Pb1 complex will neither engage with anionic membranes nor bind Pb^2+^ at Site 2. Impaired membrane binding of the Ca^2+^-sensing region of Syt1 could be a potential mechanism through which Pb^2+^ interferes with the regulation of Syt1 function by Ca^2+^ and disrupts the evoked release of neurotransmitters.^33-40^ By the same token, Pb^2+^-driven partial membrane association of Syt1 could explain how Pb^2+^ induces sporadic release of certain neurotransmitters.^37^

## CONCLUSION

Identification of oxygen-rich sites that Pb^2+^ could potentially target is essential for understanding how Pb^2+^ interferes with the function of Ca^2+^-dependent signalling proteins. Given low bioavailability of Pb^2+^, distinguishing and isolating high-from low-affinity protein sites is essential. We demonstrated using two Ca^2+^-sensing C2 domains of Syt1, that Pb^2+^-chelating pH buffer Bis-Tris provides the means to achieve such isolation. Pb^2+^ in Bis-Tris populates only one site per C2 domain. The implication for Syt1 is that two of its Ca^2+^-binding sites (out of total five) are likely to be targeted by Pb^2+^ in cellular milieu that contains natural chelators. The effect of Bis-Tris on the metal-ion dependent membrane interactions of C2 domains revealed their differential response to Pb^2+^. Our results suggest that C2B rather than C2A mediates the Pb^2+^-driven association of Syt1 with anionic membranes.

Since protein-metal ion interactions are at the core of many biological processes, extensive biochemical analysis of the corresponding binding equilibria is necessary. The use of non-chelating pH buffers is intuitively preferred, to avoid possible complications presented by metal ion chelation. This work provides a different perspective. If used in combination with non-chelating buffers, chelating pH agents, such as Bis-Tris, could potentially provide valuable mechanistic information that could otherwise be overlooked.

## Author Contributions

T.I.I. and S.K. designed the study. S.K. conducted the experiments and analyzed the data. T.I.I. and S.K. wrote the manuscript.

## Acknowledgement

This work was supported in part by NSF CHE-1905116, NIH R01GM108998, and Welch A-1784, all to T.I.I.

## REFERENCES

1. S. Caito, B. A. Lopes Ana Carolina, M. B. Paoliello Monica and M. Aschner, in Lead – Its Effects on Environment and Health, 2017, vol. 17.

2. T. I. Lidsky and J. S. Schneider, Lead neurotoxicity in children: basic mechanisms and clinical correlates, Brain, 2003, 126, 5–19.

3. D. C. Bellinger, Very low lead exposures and children’s neurodevelopment, Current opinion in pediatrics, 2008, 20, 172–177.

4. R. Gorkhali, K. Huang, M. Kirberger and J. J. Yang, Defining potential roles of Pb^2+^ in neurotoxicity from a calciomics approach, Metallomics, 2016, 8, 563–578.

5. M. Kirberger, H. C. Wong, J. Jiang and J. J. Yang, Metal toxicity and opportunistic binding of Pb^2+^ in proteins, J. Inorg. Biochem., 2013, 125, 40–49.

6. M. Geppert, Y. Goda, R. E. Hammer, C. Li, T. W. Rosahl, C. F. Stevens and T. C. Sudhof, Synaptotagmin I: a major Ca^2+^ sensor for transmitter release at a central synapse, Cell, 1994, 79, 717–727.

7. C. M. Bouton, L. P. Frelin, C. E. Forde, H. Arnold Godwin and J. Pevsner, Synaptotagmin I is a molecular target for lead, J. Neurochem., 2001, 76, 1724–1735.

8. S. Katti, B. Her, A. K. Srivastava, A. B. Taylor, S. W. Lockless and T. I. Igumenova, High affinity interactions of Pb^2+^ with synaptotagmin I, Metallomics, 2018, 10, 1211–1222.

9. K. H. Scheller, T. H. Abel, P. E. Polanyi, P. K. Wenk, B. E. Fischer and H. Sigel, Metal ion/buffer interactions. Stability of binary and ternary complexes containing 2-[bis(2-hydroxyethyl)amino]-2(hydroxymethyl)-1,3-propanediol (Bistris) and adenosine 5’-triphosphate (ATP), Eur. J. Biochem., 1980, 107, 455–466.

10. C. M. H. Ferreira, I. S. S. Pinto, E. V. Soares and H. M. V. M. Soares, (Un)suitability of the use of pH buffers in biological, biochemical and environmental studies and their interaction with metal ions – a review, RSC Advances, 2015, 5, 30989–31003.

11. K. A. Morales, M. Lasagna, A. V. Gribenko, Y. Yoon, G. D. Reinhart, J. C. Lee, W. Cho, P. Li and T. I. Igumenova, Pb^2+^ as modulator of protein-membrane interactions, J. Am. Chem. Soc., 2011, 133, 10599–10611.

12. D. Long and D. Yang, Buffer interference with protein dynamics: a case study on human liver fatty acid binding protein, Biophys. J., 2009, 96, 1482–1488.

13. A. B. Seven, K. D. Brewer, L. Shi, Q. X. Jiang and J. Rizo, Prevalent mechanism of membrane bridging by synaptotagmin-1, Proc. Natl. Acad. Sci. U. S. A., 2013, 110, E3243–3252.

14. R. Fernandez-Chacon, A. Konigstorfer, S. H. Gerber, J. Garcia, M. F. Matos, C. F. Stevens, N. Brose, J. Rizo, C. Rosenmund and T. C. Sudhof, Synaptotagmin I functions as a calcium regulator of release probability, Nature, 2001, 410, 41–49.

15. X. Zhang, J. Rizo and T. C. Sudhof, Mechanism of phospholipid binding by the C2A-domain of synaptotagmin I, Biochemistry, 1998, 37, 12395–12403.

16. S. Katti, S. B. Nyenhuis, B. Her, D. S. Cafiso and T. I. Igumenova, Partial metal ion saturation of C2 domains primes Syt1-membrane interactions, bioRxiv, 2019, DOI: 10.1101/810010, 810010.

17. M. Hothorn, I. D’Angelo, J. A. Marquez, S. Greiner and K. Scheffzek, The invertase inhibitor Nt-CIF from tobacco: a highly thermostable four-helix bundle with an unusual N-terminal extension, J. Mol. Biol., 2004, 335, 987–995.

18. S. Y. Park, W. R. Lee, S. C. Lee, M. H. Kwon, Y. S. Kim and J. S. Kim, Crystal structure of single-domain VL of an anti-DNA binding antibody 3D8 scFv and its active site revealed by complex structures of a small molecule and metals, Proteins, 2008, 71, 2091–2096.

19. H. T. Do, H. Li, G. Chreifi, T. L. Poulos and R. B. Silverman, Optimization of Blood-Brain Barrier Permeability with Potent and Selective Human Neuronal Nitric Oxide Synthase Inhibitors Having a 2-Aminopyridine Scaffold, J. Med. Chem., 2019, 62, 2690–2707.

20. J. A. Corbin, J. H. Evans, K. E. Landgraf and J. J. Falke, Mechanism of specific membrane targeting by C2 domains: localized pools of target lipids enhance Ca^2+^ affinity, Biochemistry, 2007, 46, 4322–4336.

21. K. Duncan, Metallothioneins and Related Chelators. Metal Ions in Life Sciences vol. 5. edited by Astrid Sigel, Helmut Sigel and Roland K. O. Sigel, 2009, 48, 7966–7967.

22. J. Bai, W. C. Tucker and E. R. Chapman, PIP2 increases the speed of response of synaptotagmin and steers its membrane-penetration activity toward the plasma membrane, Nat. Struct. Mol. Biol., 2004, 11, 36–44.

23. A. Perez-Lara, A. Thapa, S. B. Nyenhuis, D. A. Nyenhuis, P. Halder, M. Tietzel, K. Tittmann, D. S. Cafiso and R. Jahn, PtdInsP2 and PtdSer cooperate to trap synaptotagmin-1 to the plasma membrane in the presence of calcium, Elife, 2016, 5.

24. M. Xue, C. Ma, T. K. Craig, C. Rosenmund and J. Rizo, The Janus-faced nature of the C_2_B domain is fundamental for synaptotagmin-1 function, Nat. Struct. Mol. Biol., 2008, 15, 1160–1168.

25. G. van den Bogaart, K. Meyenberg, U. Diederichsen and R. Jahn, Phosphatidylinositol 4,5-bisphosphate increases Ca^2+^ affinity of synaptotagmin-1 by 40-fold, J. Biol. Chem., 2012, 287, 16447–16453.

26. S. Katti, S. B. Nyenhuis, B. Her, A. K. Srivastava, A. B. Taylor, P. J. Hart, D. S. Cafiso and T. I. Igumenova, Non-Native Metal Ion Reveals the Role of Electrostatics in Synaptotagmin 1-Membrane Interactions, Biochemistry, 2017, 56, 3283–3295.

27. M. S. Perin, N. Brose, R. Jahn and T. C. Sudhof, Domain structure of synaptotagmin (p65), J. Biol. Chem., 1991, 266, 623–629.

28. D. Arac, X. Chen, H. A. Khant, J. Ubach, S. J. Ludtke, M. Kikkawa, A. E. Johnson, W. Chiu, T. C. Sudhof and J. Rizo, Close membrane-membrane proximity induced by Ca(2+)-dependent multivalent binding of synaptotagmin-1 to phospholipids, Nat. Struct. Mol. Biol., 2006, 13, 209–217.

29. D. Z. Herrick, W. Kuo, H. Huang, C. D. Schwieters, J. F. Ellena and D. S. Cafiso, Solution and membrane-bound conformations of the tandem C2A and C2B domains of synaptotagmin 1: Evidence for bilayer bridging, J. Mol. Biol., 2009, 390, 913–923.

30. M. Ahamed and M. K. Siddiqui, Low level lead exposure and oxidative stress: current opinions, Clin. Chim. Acta, 2007, 383, 57–64.

31. W. Gordy and W. J. O. Thomas, Electronegativities of the Elements, 1956, 24, 439–444.

32. T. Dudev, C. Grauffel and C. Lim, How Pb^2+^ Binds and Modulates Properties of Ca^2+^-Signaling Proteins, Inorg. Chem., 2018, 57, 14798–14809.

33. R. S. Manalis and G. P. Cooper, Letter: Presynaptic and postsynaptic effects of lead at the frog neuromuscular junction, Nature, 1973, 243, 354–356.

34. P. T. Carroll, E. K. Silbergeld and A. M. Goldberg, Alteration of central cholinergic function by chronic lead acetate exposure, Biochem. Pharmacol., 1977, 26, 397–402.

35. D. J. Minnema, R. D. Greenland and I. A. Michaelson, Effect of in vitro inorganic lead on dopamine release from superfused rat striatal synaptosomes, Toxicol. Appl. Pharmacol., 1986, 84, 400–411.

36. D. J. Minnema, I. A. Michaelson and G. P. Cooper, Calcium efflux and neurotransmitter release from rat hippocampal synaptosomes exposed to lead, Toxicol. Appl. Pharmacol., 1988, 92, 351–357.

37. L. Struzynska and U. Rafalowska, The effect of lead on dopamine, GABA and histidine spontaneous and KCl-dependent releases from rat brain synaptosomes, Acta Neurobiol. Exp. (Wars.), 1994, 54, 201–207.

38. S. M. Lasley and M. E. Gilbert, Presynaptic glutamatergic function in dentate gyrus in vivo is diminished by chronic exposure to inorganic lead, Brain Res., 1996, 736, 125–134.

39. M. T. Antonio and M. L. Leret, Study of the neurochemical alterations produced in discrete brain areas by perinatal low-level lead exposure, Life Sci., 2000, 67, 635–642.

40. S. M. Lasley and M. E. Gilbert, Rat hippocampal glutamate and GABA release exhibit biphasic effects as a function of chronic lead exposure level, Toxicol. Sci., 2002, 66, 139–147.

