## Supporting information for "Interference of pH buffer with Pb^2+^-peripheral domain interactions: obstacle or opportunity?"

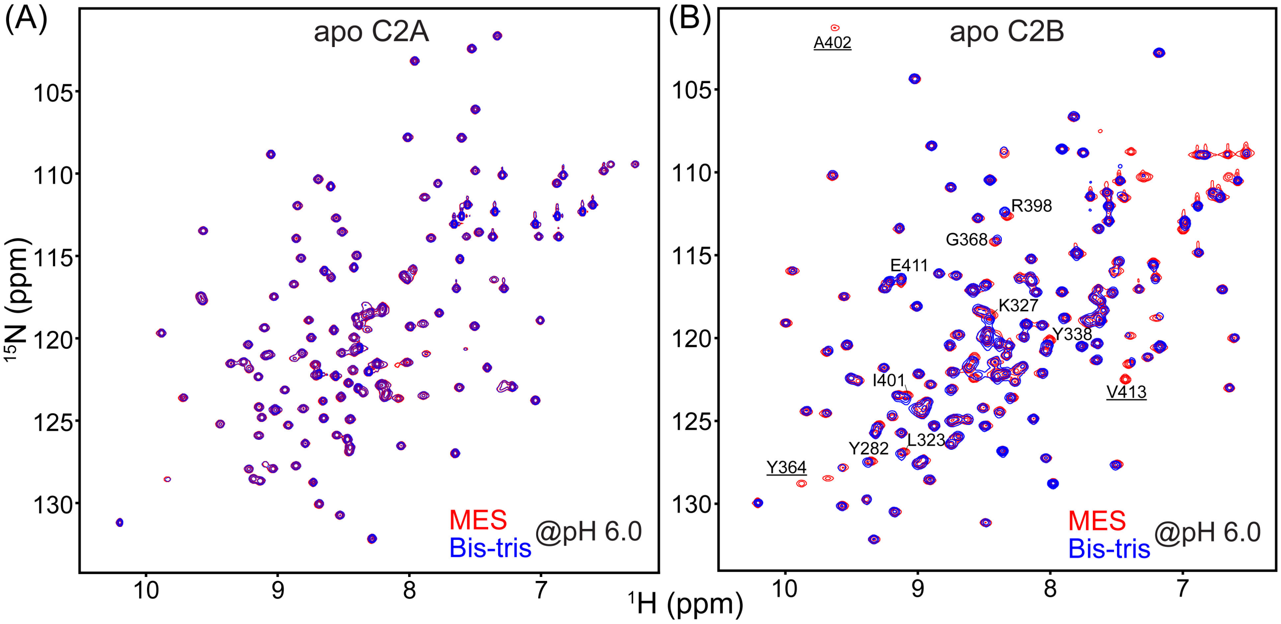


**Figure S1. C2A and C2B do not specifically interact with buffering agents.** Overlays of the [$^{15}$N-$^{1}$H] HSQC spectra of the metal-free forms of the Syt1 C2A **(A)** and C2B **(B)** domains. The C2A spectra are completely superimposable **(A)**. A subset of resonances in **(B)** shows small differences and intensity changes between the two buffers. Because of the highly basic nature of C2B and its propensity to interact with polyanionic molecules, the interacting buffer is likely to be MES. The spectra were collected at 25 °C on the Avance III NMR instruments (Bruker Biospin) operating at the magnetic field of 14.1 T/equipped with a cryoprobe (C2A); and 11.7 T/equipped with the room-temperature probe (C2B). Protein concentration in the NMR samples was 100 μM. The buffer conditions were: 20 mM MES or Bis-Tris at pH 6.0, and 150 mM KCl.

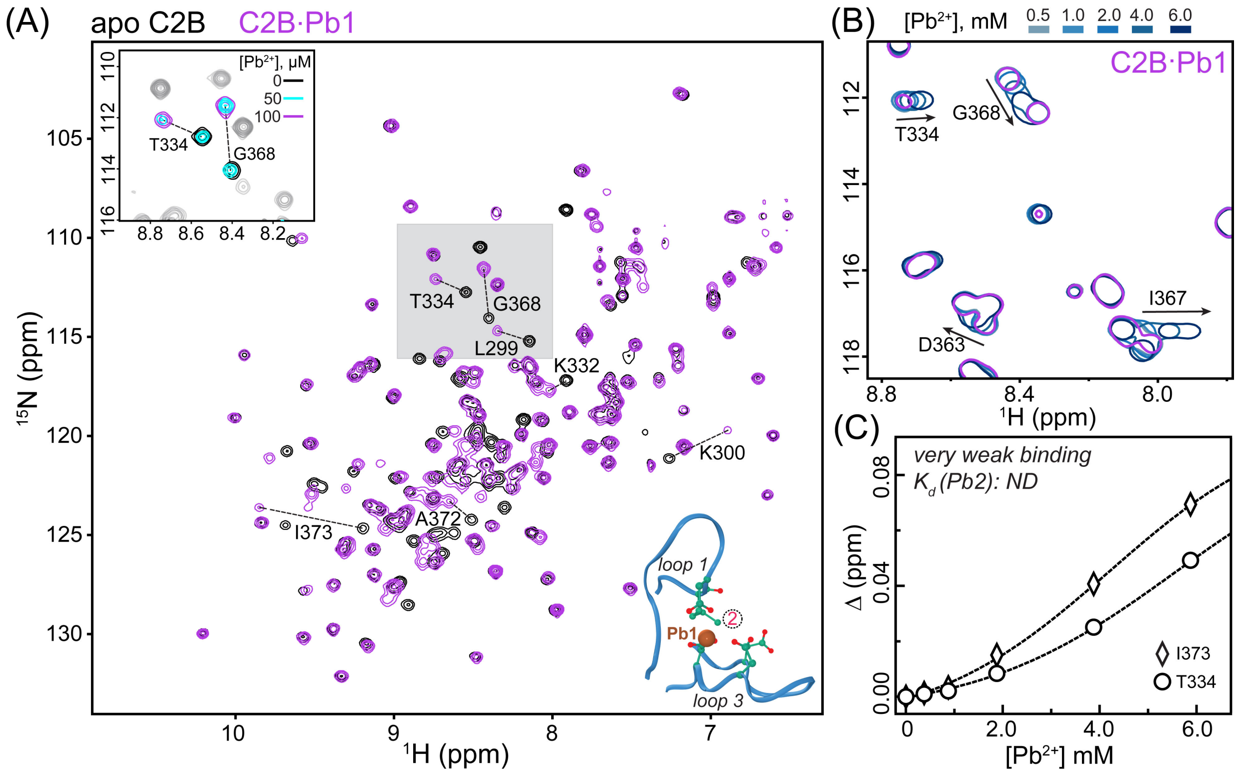


**Figure S2. Bis-Tris inhibits $Pb^{2+}$ binding to Site 2 but not Site 1 of the C2B domain. (A)** Overlay of the [$^{15}$N-$^{1}$H] HSQC spectra in the absence (apo) and presence of stoichiometric $Pb^{2+}$ (C2B:$Pb^{2+}$ 1:1, 100 µM each) in 20 mM Bis-Tris buffer at pH 6.0. Residues showing response to population of Site 1 by $Pb^{2+}$ are labeled. The spectra were collected at 25 °C on the Avance III NMR instrument (Bruker Biospin) operating at the magnetic field of 14.1 T and equipped with a cryogenically cooled probe. Top inset: expansion of the shaded spectral region with an additional $Pb^{2+}$ concentration point (50 µM, C2B:$Pb^{2+}$ 2:1, cyan) to illustrate the distinct chemical shifts of apo C2B and the C2B·Pb1 complex. Bottom inset: loop regions of the C2B domain showing the positions of Sites 1 and 2. **(B)** Overlay of the [$^{15}$N-$^{1}$H] HSQC expansions showing the chemical shift perturbations of several residues in the C2B·Pb1 complex due to $Pb^{2+}$ binding to Site 2. **(C)** Chemical shift perturbations Δ (calculated as described in ref. 8 of the main text) of two representative residues plotted as a function of $Pb^{2+}$ concentration. The non-saturatable profiles indicate that binding is extremely weak. The lines are to guide the eye.

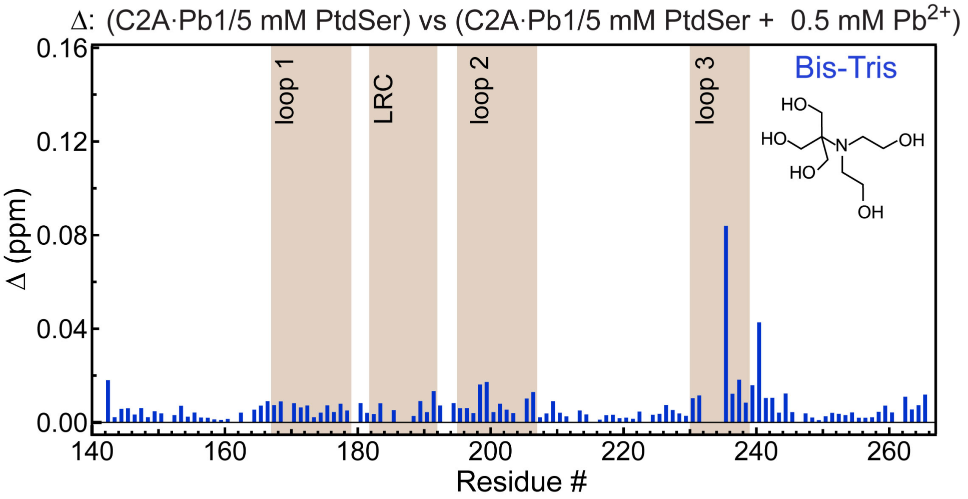


**Figure S3. PtdSer does not appreciably enhance the affinity of $Pb^{2+}$ to Site 2 of the C2A domain.** The C2A·Pb1 complex was prepared by mixing stoichiometric amounts of [U-$^{15}$N] C2A and $Pb^{2+}$ in the presence of 5 mM DPS (1,2-dihexanoyl-sn-glycero-3-phospho-L-serine, 06:0 PS). The total protein concentration was 100 µM. Addition of 0.5 mM $Pb^{2+}$ caused very weak chemical shift perturbations in the [$^{15}$N, $^{1}$H] HSQC spectra of the C2B·Pb1 complex, as shown by the plot of chemical shift perturbation Δ versus primary structure. Small perturbations around loop 3 (compared to previously reported data for $Pb^{2+}$ to Site 2, Katti et al *Metallomics*, 2018, 10, 1211-1222) indicate that $Pb^{2+}$ interactions with Site 2 remain very weak.
